# Exploring the thermal limits of malaria transmission in the western Himalaya

**DOI:** 10.1101/2022.01.31.478411

**Authors:** Farhina Mozaffer, Gautam I. Menon, Farah Ishtiaq

## Abstract

Environmental temperature is a key driver of malaria transmission dynamics. Using detailed temperature records from four sites (1800-3200m) in the western Himalaya, we model how temperature regulates parasite development rate (the inverse of the extrinsic incubation period, EIP) in the wild. Using a Briére parametrization of the EIP, combined with Bayesian parameter inference, we study the thermal limits of transmission for avian (*P. relictum*) and human *Plasmodium* parasites (*P. vivax* and *P. falciparum*) as well as for two malaria-like avian parasites, *Haemoproteus* and *Leucocytozoon*. We demonstrate that temperature conditions can substantially alter the incubation period of parasites at high elevation sites (2600-3200m) leading to restricted parasite development or long transmission windows. We then compare estimates of EIP based on measures of mean temperature versus hourly temperatures to show that EIP days vary in cold versus warm environments. We found that human *Plasmodium* parasites experience a limited transmission window at 2600m. In contrast, for avian *Plasmodium* transmission was not possible between September to March at 2600m. In addition, temperature conditions suitable for both *Haemoproteus* and *Leucocytozoon* transmission were obtained from June to August and in April, at 2600m. Finally, we use temperature projections from a suite of climate models to predict that by 2040, high elevation sites (~ 2600 m) will have a temperature range conducive for malaria transmission, albeit with a limited transmission window. Our study highlights the importance of accounting for fine-scale thermal effects in the expansion of the range of the malaria parasite with global climate change.

---

Changes in climate are shifting the geographic ranges of species (e.g., birds; Chen *et al*. 2011, Freeman *et al*. 2018). These have a significant influence on parasite transmission dynamics, either by exposing immunologically naïve hosts to longer transmission seasons or bridging novel host-parasite interactions (Patz *et al*. 2001). Among vector-borne pathogens, malaria parasites have remained the most virulent group, with high sensitivity to climatic factors which continue to threaten both human and many wild animal populations (van Riper *et al*. 1999).

Temperature is the key environmental driver influencing the transmission dynamics and distribution of malaria parasites. The rate of malaria parasite transmission and intensity of infection is strongly determined by the extrinsic incubation period (EIP: also known as the duration of sporogony), the time it takes for a parasite to develop within a mosquito and become transmissible (Ohm *et al*. 2018). Therefore, the EIP determines the parasite development rate in the midgut after many replication cycles before it migrates as a sporozoite (infective stage) in the salivary glands of an arthropod vector. The development rate of a parasite depends on host, parasite, and environmental conditions. These conditions must be conducive for transmission of the parasite. For example, the EIP of human *Plasmodium* is dependent on temperature and the parasite generally takes 8–14 days to develop, so if adult *Anopheles* mosquitoes die before or within a 12-day period, they are unlikely to contribute to parasite transmission (Killeen *et al*. 2000, Paijmans *et al*. 2009, Ohm *et al*. 2018). In addition, temperature plays a central role in regulating mosquito population dynamics, age-structure in a population, life-history traits, fitness and phenology of vectors and parasites, leading to complex spatial and temporal patterns of distribution (Beck-Johnson *et al*. 2013).

Most mechanistic models of human *Plasmodium* transmission are based on the degree-day model of Detinova (Detinova 1962). The Detinova model assumes a linear relationship between ambient temperature (*T*) and the parasite development rate (PDR). For example, for *Plasmodium falciparum* the following relationship is assumed to hold: EIP (in days) =111/(*T*-16), where 111 is the cumulative number of degree-days required for the parasite to complete development, *T* is the average ambient environmental temperature, and the relationship assumes that the temperature threshold below which development cannot occur is 16°C. However, although this equation is used in many studies, it fails to capture daily temperature fluctuations which could potentially alter the rate of parasite development and malaria transmission in a population. Other thermodynamic models propose a nonlinear relationship between temperature and growth or development (Paaijmans *et al*. 2009). These models can be generalised to consider temperature fluctuations which alter the length of parasite incubation period and malaria transmission rates. Therefore, epidemiological models should benefit from combining local meteorological data and diurnal temperature fluctuations to understand the biological significance of temperature in shaping parasite transmission dynamics.

Temperature is considered as the main driver for mosquito emergence and spring phenology (e.g., budburst, leaf-out, and flowering) in temperate regions (e.g., H□llfors *et al*. 2020). In the context of epidemiology of avian malaria, the overlap in seasonal emergence of vectors produces spring relapses in chronic infections (parasite phenology) and new infections in breeding host populations (Applegate 1970, Beaudoin *et al*. 1971). However, the implications of temperature variation for avian malaria parasite development across temperate regions are less understood.

The western Himalayas are a species-rich and highly seasonal ecosystem with distinct physiographic climatic conditions which drives bird migration, spring phenology and vector emergence (Barve *et al*. 2016, Ishtiaq and Barve 2018). In general, birds harbour a huge diversity of three genera of haemosporidian parasites - *Plasmodium, Haemoproteus* and *Leucocytozoon*, which are more ubiquitous and cosmopolitan (except Antarctica) (Valkinũas 2005). These parasites are transmitted by dipteran insects, e.g., mosquitoes (*Plasmodium*), biting midges (*Haemoproteus*) and black flies (*Leucocytozoon*) and have significant negative effects on the host survival and longevity (Asghar *et al*. 2015), reproductive success and body condition (Marzal *et al*. 2005). In this montane system, birds exhibit two migration strategies; species are either year-round high elevation residents (sedentary) or seasonal elevational migrants. Elevational migrants winter at low elevations or in the plains (≤ 1500 m above sea level; a.s.l.) and move to breeding grounds at higher elevations (2600□4000 m a.s.l. or even higher) during the summer season (Dixit *et al*. 2016). The elevational migrants are exposed to a large suite of parasites and vector fauna, especially in low elevations and move to high elevation breeding grounds only during the summer season (“migratory escape”, Loehle 1995). By contrast, sedentary counterparts potentially experience little or no exposure to parasites at high elevations in winters. Given that a competent vector and optimal thermal conditions are present, elevational migrants could act as ‘bridge hosts’ of parasite species and potentially increase transmission risk between wintering and breeding areas. This potentially increases the risk of infection to naïve resident birds at high elevations which might not have evolved to cope with parasite infection. In addition, the emergence of insect vectors (e.g., *Culicoides*) is driven by temperature and does not coincide with peak bird breeding season (April-May) in a high elevation environment suggesting a mismatch in phenology of vectors and avian hosts (Ishtiaq *et al*. unpubl., e.g., Gethigs *et al*. 2015). This mismatch potentially alters the degree of interaction between host and vector species, thereby influencing parasite transmission dynamics. Only if the arrival of infected birds (with infective stages in the bloodstream) coincides with the peak of vector abundance, can transmission of pathogens from migratory birds to vectors be facilitated. There are currently no studies undertaken to understand the influence of environmental factors on vector phenology and what changes in parasites’ distribution ranges are expected with climate change.

In this study we model the change in temperature and parasite transmission dynamics in four western Himalayan sites across an elevational gradient. Using fine-scale meteorological data, we explore limits of parasite transmission as a function of temperature in the western Himalayan landscape. Specifically, we ask the following questions:

i. Is parasite transmission restricted by temperature in high elevation environments?
ii. Is there spatial and temporal variation in parasite transmission dynamics?
iii. How will climate change affect the distribution of malaria and the parasite transmission window?

## MATERIALS AND METHODS

### Study area and collection of temperature data

For micro-climate data, we deployed the Thermochron iButton (Maxim Integrated Products, http://www.maxim-ic.com/) to record temperature data every hour on a round-the-clock basis at each sampling site from 2014-2015: Mandal [1800m: N30.44685° E79.27328°], Anusuya [2000-2200m: N30.47888°E79.28503°], Kanchula [2600m: N30.45913° E79.22744°] and Shokharakh [3100-3400m: N30.47860°E79.217980°; Figure 1.] in the western Himalaya, Uttarakhand state, India.

**Figure 1.**
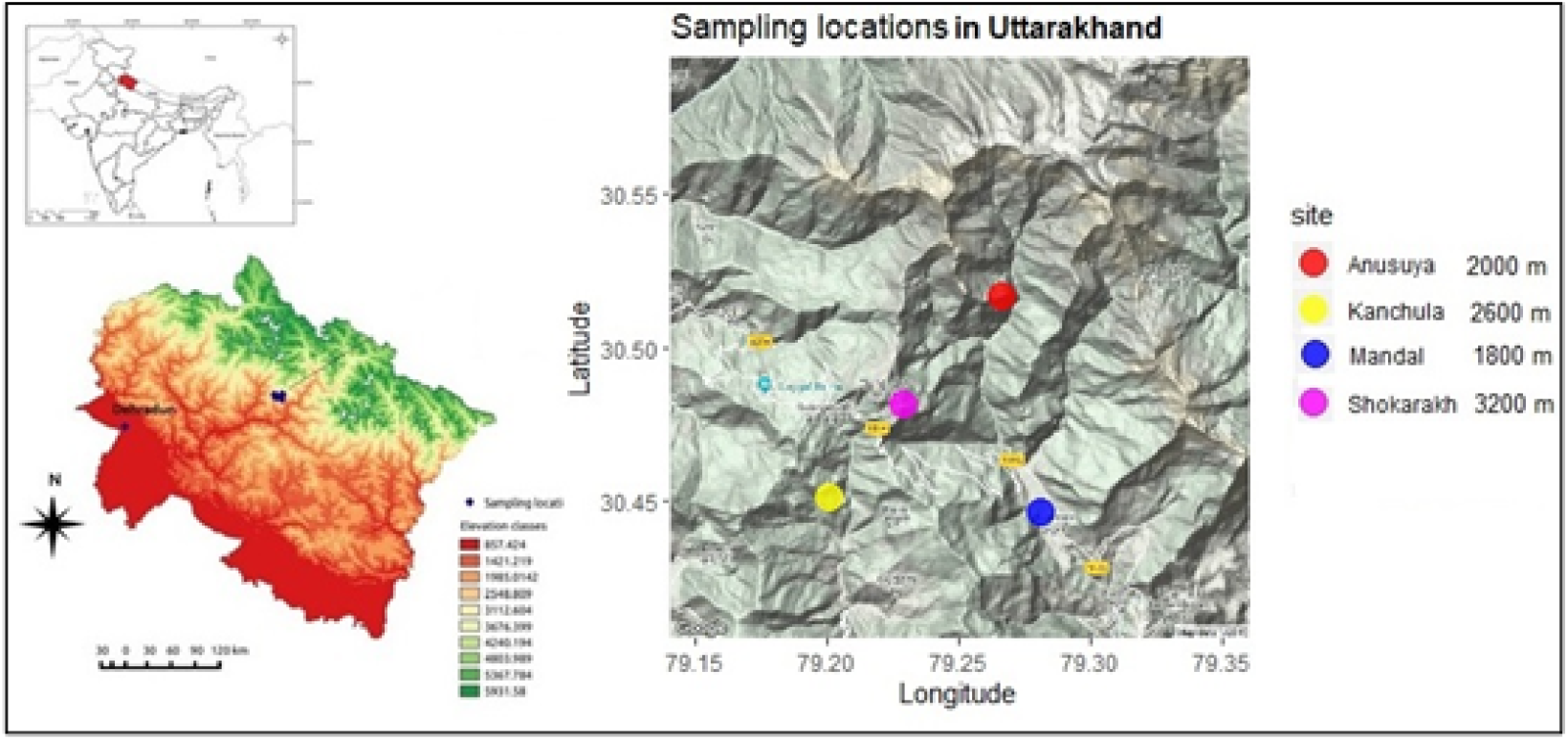
Daily temperature sampling site of the western Himalaya Mandal (1800 m), Anusuya (2000 m), Kanchula (2600 m) and Shokharakh (3200 m). We have used the Madhmaheshwar site as an alternative to Shokharakh for modelling future parasite range expansion, as explained in the main text.

### Quantifying the effects of environmental temperature on parasite development (Extrinsic Incubation Period)

The extrinsic incubation period is the reciprocal of the parasite development rate (PDR). A convenient representation of the PDR follows from the Briére equation (Briére *et al*. 1999)

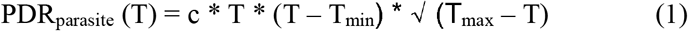

where T_min_ and T_max_ are the minimum and maximum temperatures that can sustain parasite development and c is a constant, the scaling parameter. These parameters must be estimated or fit against experimental data. This model was based on thermodynamic principles and shown to capture accurately the form of the EIP in *Plasmodium falciparum* and *Plasmodium vivax* as described previously by others (Paaijmans *et al*. 2009, Cator *et al*. 2013).

### A Bayesian approach to the calculation of the extrinsic incubation period

Given the Briére equation (Briére *et al*. 1999) we first use Bayesian methods to find the best estimates for the parameters T_min_, T_max_ and c (Gelman *et al*. 2013, Bolstad *et al*. 2016, McElreath 2016). Bayesian inference approach allows us to incorporate prior knowledge into a description of data, capturing parameter uncertainties. It also provides a consistent framework for including prior information, while accounting for uncertainty in inferred parameters.

We used thermodynamic models to estimate the influence of both mean and diurnal temperature fluctuation on malaria transmission (Paaijmans *et al*. 2009). Specifically, we selected parasite species that have been recorded in the western Himalayan birds. We extracted the temperature dependent EIP values for two human malaria parasites - *P. falciparum, P. vivax* and avian parasites in the genus *Plasmodium* (*Plasmodium relictum*), *Haemoproteus* and *Leucocytozoon* using parasite specific temperature data summarised in Suppl. Table 1. Specifically, for *Haemoproteus*, we choose the average (midpoint of the range) as only temperature ranges were available (Suppl. Table 1).

To estimate the EIP of a parasite species, two main parameters are required: the lifespan of arthropod vectors and temperature data. The two human malaria parasites - *P. falciparum, P. vivax* are transmitted by species of *Anopheles* mosquito. The avian malaria, *P. relictum*, is primarily transmitted by *Culex quinquefasciatus*. It takes the malaria parasite 56 days to develop in the mosquito at 18°C which is longer than the lifespan of the mosquitoes. At 22°C it takes only 19 days and at 30°C only 8 days (Githeko 2007). The upper limit of longevity of mosquitoes can be up to 56 days as used in previous studies (Craig *et al*. 1999, Paaijmaans *et al*. 2009) depending upon environmental conditions. We considered an upper threshold for the EIP for mosquitoes as 56 days. The vectors of *Haemoproteus* sp. are biting midges (*Culicoides* sp.) and black flies (simulids) for *Leucocytozoon*. The lifespan of biting midges is ~10 to 20 days (Sick *et al*. 2019) whereas black flies can survive for two to three weeks (Adler 2004). We thus considered the upper threshold for the EIP for both biting midges and black flies to be 20 days.

We describe our methods in more detail in subsections below:

i. **Estimation of model parameters** We estimate the parameters θ = {T_min_,T_max_, c, σ} using Bayesian inference methods (Gelman *et al*. 2013, Bolstad *et al*. 2016, McElreath 2016) implemented using the available data. The parameters T_min_, T_max_ and c have been defined earlier. The quantity σ provides the scale factor for the width of the likelihood between observed data and model. Our implementation of the inference strategy uses STAN (https://mc-stan.org/), a platform for statistical modelling and high-performance statistical computation. STAN performs Bayesian statistical inference with Markov Chain Monte Carlo (MCMC) sampling. It also provides diagnostic tools to evaluate the accuracy and convergence of the MCMC while allowing for posterior predictive checks.
ii. **Prior distribution** Bayesian approaches incorporate prior knowledge about the parameters into the model. Our choices for these distributions are summarised in (Suppl. Table 2) below for each parasite. We choose a common prior for Sigma, 1/gamma (0.0001,0.0001) and scale parameter (c), gamma (1,10), in all cases. We selected a flat prior for T_min_ and T_max_ within a defined range. We used a gamma distribution as both the scaling parameter and sigma are non-negative continuous positive values.
iii. **Likelihood** We choose a normal distribution with mean parameter μ given by the Briére equation (Briére *et al*. 1999) as the likelihood of the data and standard deviation σ. We run STAN (https://mc-stan.org/) for four chains of 1000 iterations each, discarding 500 iterations in each case for warmup. 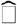, the convergence statistic reported by STAN, is close to 1 (< 1.05), indicating the 4 Markov chains are in close agreement with one another.

### Calculation of the extrinsic incubation period (EIP)

The EIP was calculated using mean daily temperature and mean hourly temperature for *P. falciparum, P. vivax, P. relictum, Leucocytozoon* and *Haemoproteus* parasites. The mean hourly temperature is used to estimate the effect of diurnal temperature fluctuation on EIP. The diurnal temperature range (DTR) variation across these four sites is the difference between maximum and minimum temperatures on each day of the month at each site. This was done for each day of the month for all four sites. Since all four sites are at different elevations, it is possible that for a few days of a month, the temperature was marginally above the minimum threshold temperature for the development of parasites. The EIP calculated using these temperatures can thus become anomalously high. We thus set the EIP to 90 days whenever the computed EIP exceeded that number. We calculated the mean EIP using both the mean daily temperature as well as mean hourly temperatures for the month with the following procedure: We calculate the EIP using the mean temperature for each day of the month. We then average this EIP over the entire month. We use a similar procedure for the mean hourly temperature. We set the upper threshold temperature to about 5°C above the maximum threshold temperature (T_max_) following Hu and Appel (2004) and Cator *et al*. (2013). The upper threshold temperature is lethal for parasite development. Once the temperature exceeds this lethal temperature the development of parasites does not occur and no estimate of EIP is possible.

### Modelling future parasite range expansion with warming climate

We used a series of climate-based models to predict the change in parasite range with temperature. First, we extracted monthly values for minimum temperature and maximum temperature for the year 2021-2040 for four Himalayan sites. We used Madhmaheshwar as a nearby site (30°38’13”N 79°12’58”E) as an alternative to Shokharakh due to non-availability of data.

We used data from eight global climate models available on WorldClim (www.worldclim.org). These global climate models (GCM) are BCC-CSM2-MR, CNRM-CM6-1, CNRM-ESM2-1, CanESM5, IPSL-CM6A-LR, MIROC-ES2L, MIROC6, MRI-ESM2-0. We extracted temperature data for the year 2021-2040 at a spatial resolution of 2.5 minutes to estimate the effect of future climate change on malaria transmission. A set of scenarios have been chosen to provide a range of distinct end of century climate change outcomes by the energy modelling community, which mainly deals with greenhouse gas emission scenarios driven by different socioeconomic assumptions. We considered the shared socioeconomic pathway SSP2-4.5, which provides one scenario for global emissions, consistent with certain assumptions about how socio-economic trends might evolve in the future. We selected SSP2-4.5 because it represents a “middle of the road” scenario i.e., the world follows a path in which social, economic, and technological trends do not shift markedly from historical patterns.

To extract the minimum temperature and maximum temperature for the year 2021-2040 values, we use the GIS software QGIS (https://www.qgis.org). We compute the average of predicted monthly mean temperature across all eight GCM models.

### Comparison of EIP calculated using local meteorological data and WorldClim data

The temperature collected from data loggers is the real temperature experienced by the malaria vectors in the field. Therefore, data from loggers can be expected to provide more precise and local results as compared to the weather station temperature data. To understand variation in temperature modelling based on the local temperature records and WorldClim (www.worldclim.org) data, we calculated the EIP using mean temperature collected from both datasets from May to April 2014 to 2015.

## Results

We estimated the extrinsic incubation temperature of three *Plasmodium* and two malaria-like parasites-*Haemoproteus* and *Leucocytzoon* using one-year field-collected data across four sites in the western Himalaya. Mean daily temperatures were warmest (18°C) for the low elevation site at 1800m (Mandal), followed by the mid elevation site (13°C). The two high elevation sites at 2600m (Kanchula) and 3200m (Shokharakh) had the coolest temperatures, 10°C and 8°C, respectively (as shown in Figure 2).

**Figure 2.**
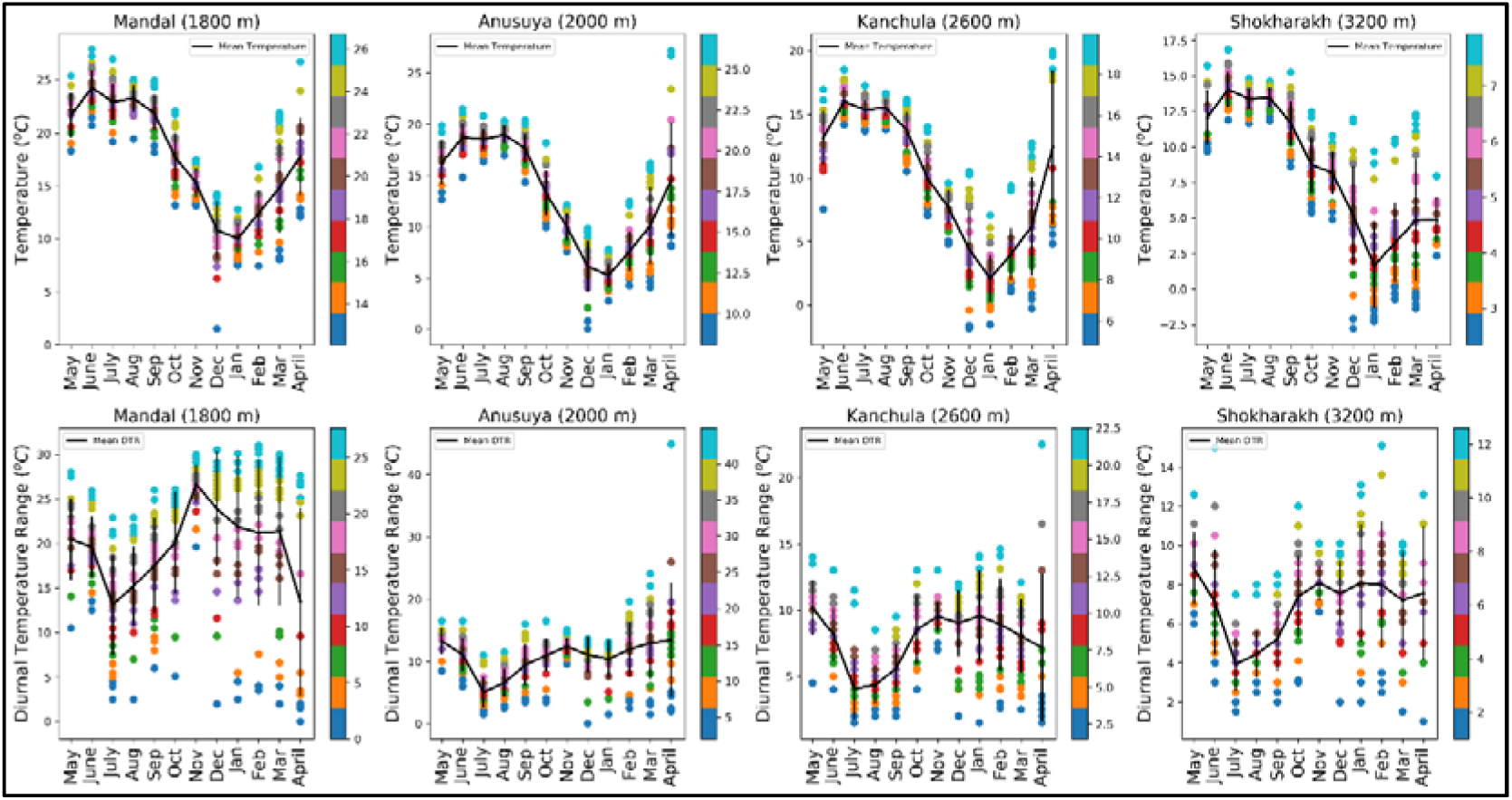
Daily temperature records at an hour interval across four sites in the western Himalaya. Mean temperature (upper panel) and DTR (Diurnal temperature range) (Lower panel) for all four sites. Each point shown in colour represents a value of daily temperature (in ^O^C, right hand bar) from May 2014 to April 2015 of that site. The solid lines represent average temperature. The black vertical bar represents standard deviation.

### Explicit formulae for the Extrinsic incubation period

The EIP of each parasite, in the Briere parametrization, requires three parameters to be specified. The distributions of these parameters must be obtained using prior knowledge and the available data, together with statistical methods. We obtained parameter values describing the EIP using the Bayesian methods described above, for *P. relictum, Haemoproteus* and *Leucocytozoon*. The parameter values obtained together with the appropriate confidence interval are shown in Table 1. We display our results in Figure. 3, together with those derived earlier for *P. falciparum* and *P. vivax* by Cator *et al*. (2013).

**Figure 3:**
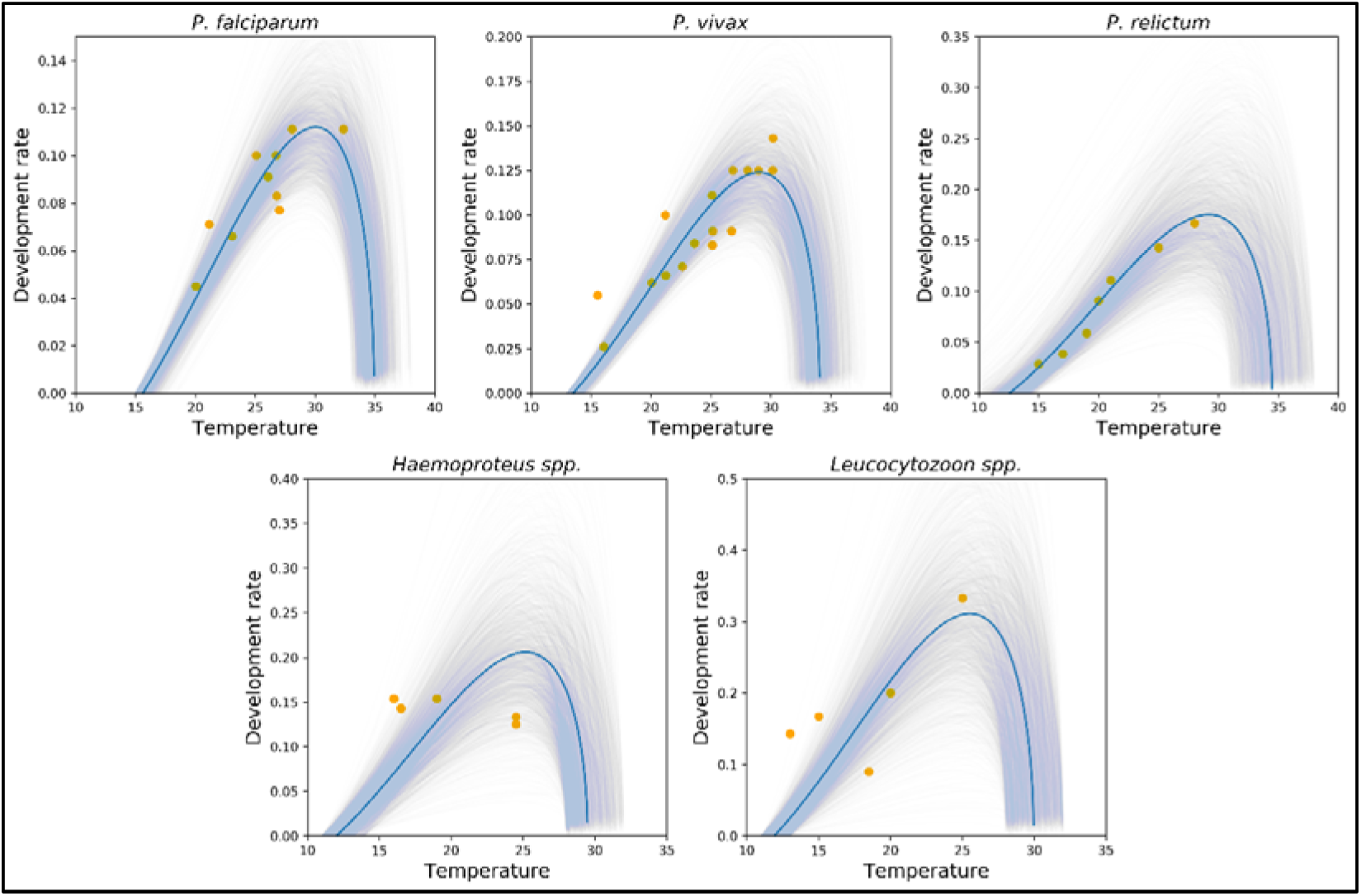
Relationship between temperature and parasite development rate for P. falciparum, P. vivax, P. relictum, Haemoproteus and Leucocytozoon obtained using Bayesian modelling as described in the text. The available empirical data (circle), summarised in the Suppl. Table 1, is fitted to the Briére equation. The prior distributions for all the estimated parameters are provided in Suppl. Table 2. The solid line is the fit to the empirical data. The bands show the uncertainty (confidence interval) in inferred parameters.

**Figure 4:**
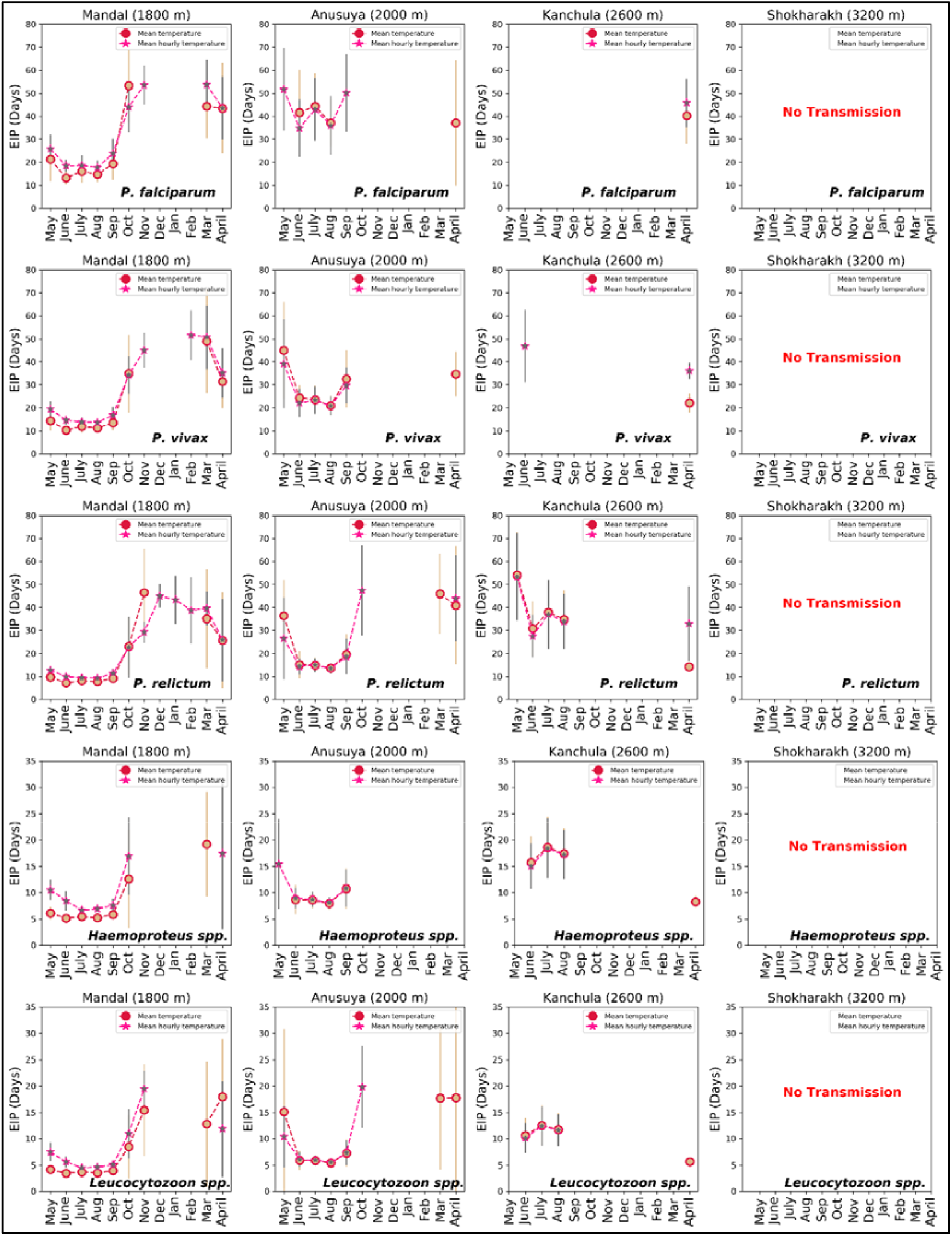
Extrinsic incubation period (EIP) in days of P. falciparum, P. vivax, P. relictum, Haemoproteus and Leucocytozoon for four sites of the western Himalaya. These are calculated using mean temperature (circle) and diurnal temperature range (DTR) (star) from May 2014 to April 2015. Bar represents the standard deviation.

**Table 1.**
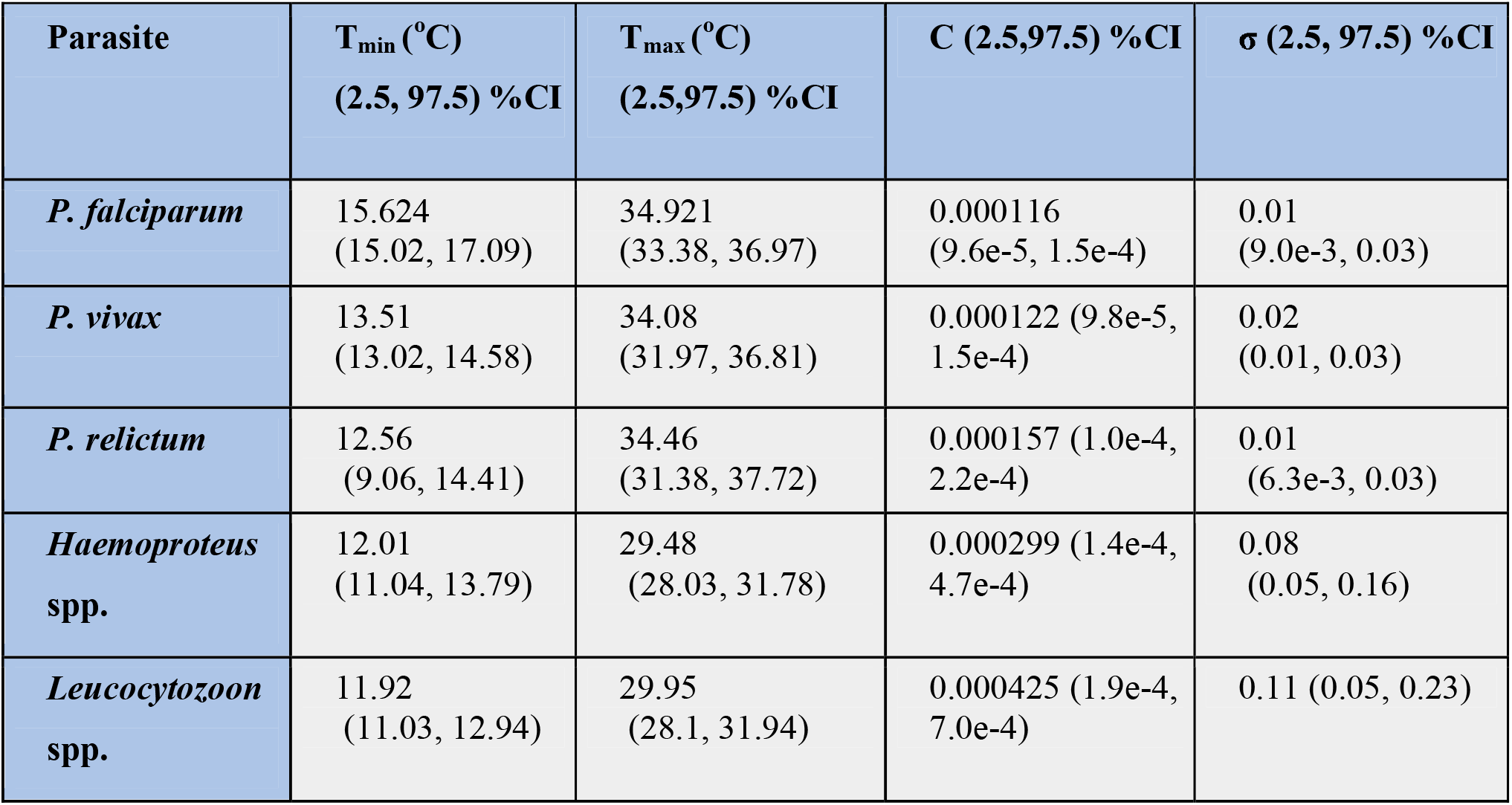
Estimated parameter values of minimum and maximum temperature, scaling parameter c and sigma using Bayesian method for all parasites. % CI is the confidence interval.

| Parasite | $T_{\min}$ (°C)<br>(2.5, 97.5) %CI | $T_{\max}$ (°C)<br>(2.5, 97.5) %CI | C (2.5, 97.5) %CI | $\sigma$ (2.5, 97.5) %CI |
| --- | --- | --- | --- | --- |
| *P. falciparum* | 15.624<br>(15.02, 17.09) | 34.921<br>(33.38, 36.97) | 0.000116<br>(9.6e-5, 1.5e-4) | 0.01<br>(9.0e-3, 0.03) |
| *P. vivax* | 13.51<br>(13.02, 14.58) | 34.08<br>(31.97, 36.81) | 0.000122 (9.8e-5,<br>1.5e-4) | 0.02<br>(0.01, 0.03) |
| *P. relictum* | 12.56<br>(9.06, 14.41) | 34.46<br>(31.38, 37.72) | 0.000157 (1.0e-4,<br>2.2e-4) | 0.01<br>(6.3e-3, 0.03) |
| *Haemoproteus*<br>spp. | 12.01<br>(11.04, 13.79) | 29.48<br>(28.03, 31.78) | 0.000299 (1.4e-4,<br>4.7e-4) | 0.08<br>(0.05, 0.16) |
| *Leucocytozoon*<br>spp. | 11.92<br>(11.03, 12.94) | 29.95<br>(28.1, 31.94) | 0.000425 (1.9e-4,<br>7.0e-4) | 0.11 (0.05, 0.23) |

For *P. falciparum* and *P. vivax*, we rederived the forms obtained by Cator *et al*. (2013) through these Bayesian methods. The work of Cator *et al*. (2013) used an alternative method involving simple curve-fitting using SPSS (IBM SPSS v20). We find exact consistency with these earlier results, demonstrating the validity of our numerical methods. Our results extrapolate from the limited data available, together with non-informative estimates of the relevant priors, to obtain what appear to be reasonable fits to the data, together with uncertainty bounds.

### Effects of environmental temperature on parasite development (Extrinsic Incubation Period)

Using the thermodynamic parasite development model, we estimated the effect of using the mean daily temperature and mean hourly temperature on the calculation of the EIP for each site (1800-3200m). Our methods described the rate of parasite development for *Plasmodium* species (two human and one avian *Plasmodium*), *Haemoproteus*, and *Leucocytozoon* parasites (Figure 3) at varying temperatures. For high elevation sites (3200m), the EIP estimates for all parasites using both measures suggested that transmission was not possible throughout the year. Below, we describe where results from both measures coincide, for other elevations and parasites, as well as where they differ in detail. We find that both measures, based on either the diurnal temperature range or the mean temperature, very largely give similar results for the periods of the year where the EIP is within the allowable range for transmission. However, for specific months, it is possible that transmission is possible within one scheme but not within the other.

For the 2600m site, there was no parasite transmission predicted using the EIP based on both temperature measures for *P. falciparum* and *P. vivax* from May to March. However, in June, the mean temperature range predicted no transmission of *P. vivax* whereas the EIP days of 46.9±15.71 using the diurnal temperature range. In April, the EIP days for *P. vivax* were longer by 13.95 days using the diurnal temperature range (EIP days: 36.08±3.5) than the mean temperature (EIP days: 22.13±4.04). The EIP days for *P. falciparum* were 5.62 days shorter using mean temperature measures (EIP days:40.29± 12.11) than the diurnal temperature (EIP days: 45.91±10.64) in April.

For avian *Plasmodium relictum*, using both mean temperature and diurnal temperature range, September to March was predicted as no transmission period and May to August and April as predicted transmission, at 2600m. In April, the EIP was 18.7 days longer using the diurnal temperature range (EIP days:32.95±16.29) than the mean temperature range (EIP days:14.25±2.25). For two malaria-like avian parasites, *Haemoproteus* and *Leucocytozoon,* September to March were predicted as no transmission period and June to August was predicted as the transmission period (< 1 day difference in EIP estimates) using both temperature measures. However, diurnal temperature range predicted no parasite transmission in April whereas mean temperature range estimated the EIP days of 8.27±1.1 and 5.62±0.75, for *Haemoproteus* and *Leucocytozoon*, respectively.

For the 2000m site (Anusuya), October to March was predicted to have no transmission period for *P. falciparum* and *P. vivax* using both temperature measures. In April, using the mean temperature predicted EIP days of 37.12±27.34 for *P. falciparum* and 34.73±9.66 for *P. vivax*, however no transmission was obtained using the diurnal temperature range. Using the mean temperature, there was no *P. falciparum* transmission predicted for May and September. However, the diurnal temperature range approach predicted EIP days of 51.7±17.94 in May and 50.7±17.02 in September.

Among avian parasites, *P. relictum* transmission window was predicted from May to September using both temperature measures. However, there was no *P. relictum* transmission predicted in October using mean temperatures while the diurnal temperature range method estimated EIP days of (47.4±19.46). In March, use of the diurnal temperature range predicted no *P. relictum* transmission whereas the mean temperature range measure suggested EIP days of 46.03±17.43. In April, both temperature measures predicted *P. relictum* transmission, however, EIP days using diurnal temperature range were 3.06 longer than for the mean temperature. For two malaria-like parasites, *Haemoproteus* and *Leucocytozoon*, transmission was predicted from June to September using both temperature measures. There was no *Haemoproteus* transmission predicted for October to April using both temperature measures. In May, *Haemoproteus* showed no transmission using mean temperature whereas, using the diurnal temperature, transmission was predicted with showed EIP days of 15.44±8.54. However, *Leucocytozoon* transmission was not predicted from November to February, while only the mean temperature supported EIP days of 17.71±13.56 in March and April. The diurnal temperature range method indicated transmission in October with EIP days of 19.81±7.8.

For the lowest elevation site (1800m), *P. falciparum* and *P. vivax* showed no transmission from December to February using both measures, the DTR and the mean temperature. An exception was for P. *vivax* where the use of the diurnal temperature range showed an EIP of 51.6±10.72 days in the month of February. The effect of daily temperature variations on *P. falciparum* and *P. vivax* predicted a transmission window from May to October, March, and April. In November, there was no transmission using the mean temperature for both *P. falciparum* and *P. vivax*. The use of the diurnal temperature range predicted transmission for both parasites in that month, with EIP more than 45 days in both cases.

In contrast, for *P. relictum*, use of the diurnal temperature range suggested transmission throughout the year. However, the use of the mean temperature showed no transmission across the months December to February. For the two malaria-like parasites, transmission months ranged from May to October for *Haemoproteus* and May to November, as well as April, for *Leucocytozoon*. The exceptions were: using the mean temperature led to transmission in March while using the diurnal temperature range led to transmission in April for *Haemoproteus*. In March, there was no transmission predicted using the diurnal temperature range while mean temperature range predicted EIP days of 12.79±11.96 for *Leucocytozoon*.

Our comparisons of mean temperature collected using experimental logger data and WorldClim data from 2014-2015 showed threshold temperature not supporting parasite transmission at the high elevation site (3200m) throughout the year (Suppl. Fig. S1). However, at the 2600 m site, *P. vivax* transmission was predicted from May to September and April whereas experimental data suggested only in April. For avian parasites, *P. relictum, Haemoproteus* and *Leucocytozoon* the window in which transmission is predicted is largely similar using these two source datasets, albeit leading to longer EIP days with experimental data, with the only exception for the month of April where this systematics are reversed. This larger pattern was reversed at 1800 m, with shorter EIP days with experimental data as compared to using WorldClim data, with the only exception again being for the month of April. The 2000m site showed closely similar transmission patterns for all parasites using both datasets, again except for the months of March and April, for *P. relictum* and *Leucocytozoon*.

### Predicting parasite range expansion

Using computed monthly mean temperatures extracted from global climate projections (2021-2040) and our field data collected from 2014-2015, we compared the EIP for avian as well as the human malaria parasites across the four Himalayan sites (Figure 5). The main point was that for virtually all parasites, climate change scenarios lead to an expansion of the transmission period in which parasite survival is guaranteed as well as the lower EIP. The only exception was the low elevation site, Mandal (1800m) where a decrease in average temperature is predicted leading to an increase in EIP. The effects of climate change are known to be inhomogeneous, in general, even as there is a secular increase in overall mean temperatures. This counterexample to the general trend supports that observation.

**Figure 5:**
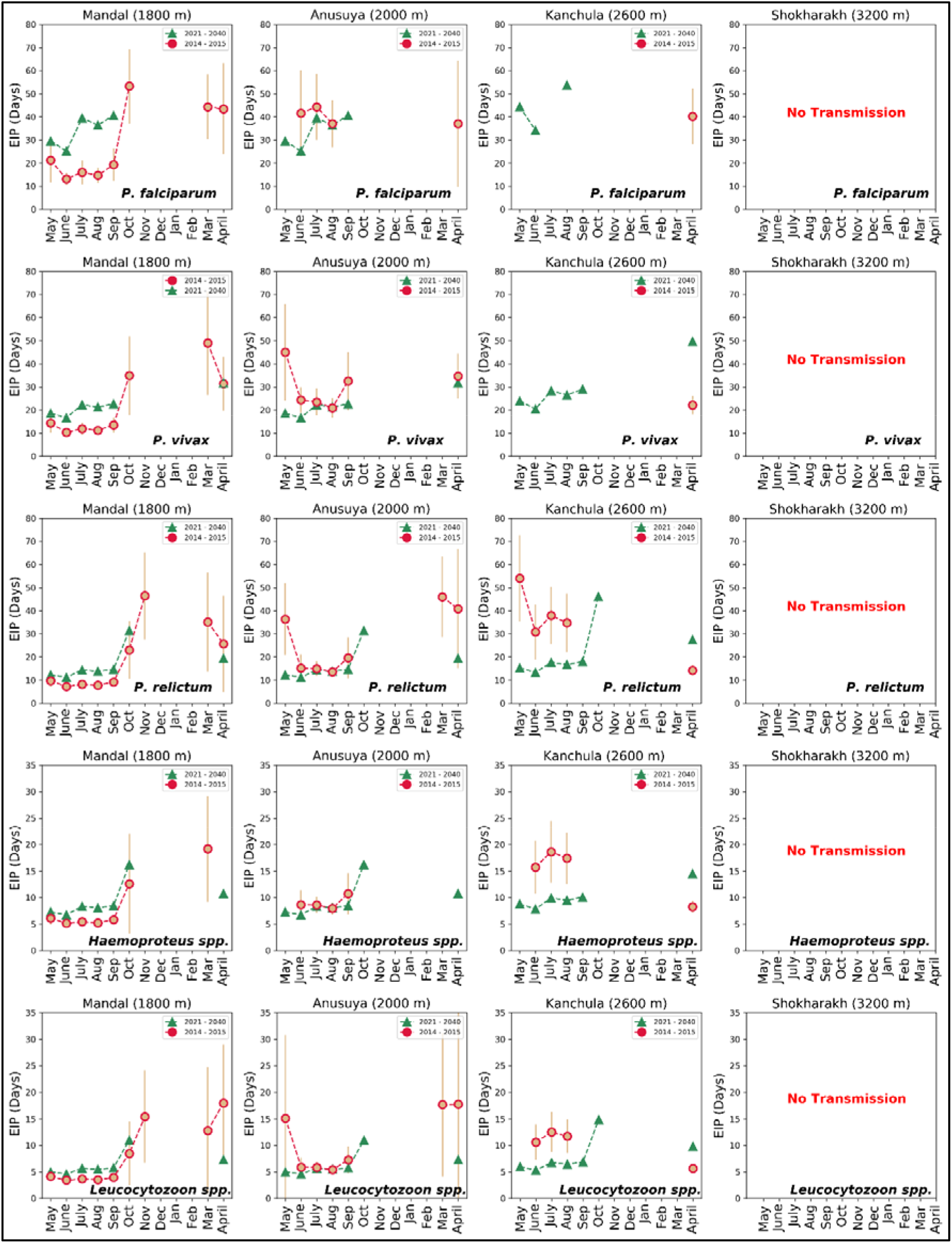
Comparison of extrinsic incubation period (EIP) in days of P. falciparum, P. vivax, P. relictum, Haemoproteus and Leucocytozoon for four sites of the western Himalaya. using mean temperature data (May 2014 to April 2015, circle) and future climate data (2021 to 2040, triangle). Bar represents the SD.

### Comparison of EIP using historical and future climate data

To understand the effect of climate change with the historical and future climate temperature on the EIP, we have used average temperature for the year 2000-2018 (historical temperature data) and average temperature of 2021-2040 (future temperature data), using data from WorldClim (www.worldclim.org). The EIP calculated using both these data is shown in Figure 6. The effects of global warming are clearly seen, i.e., with high temperature for the year 2021-2040, the EIP is shorter as compared to the lower temperatures across the period 2000-2018.

**Figure 6:**
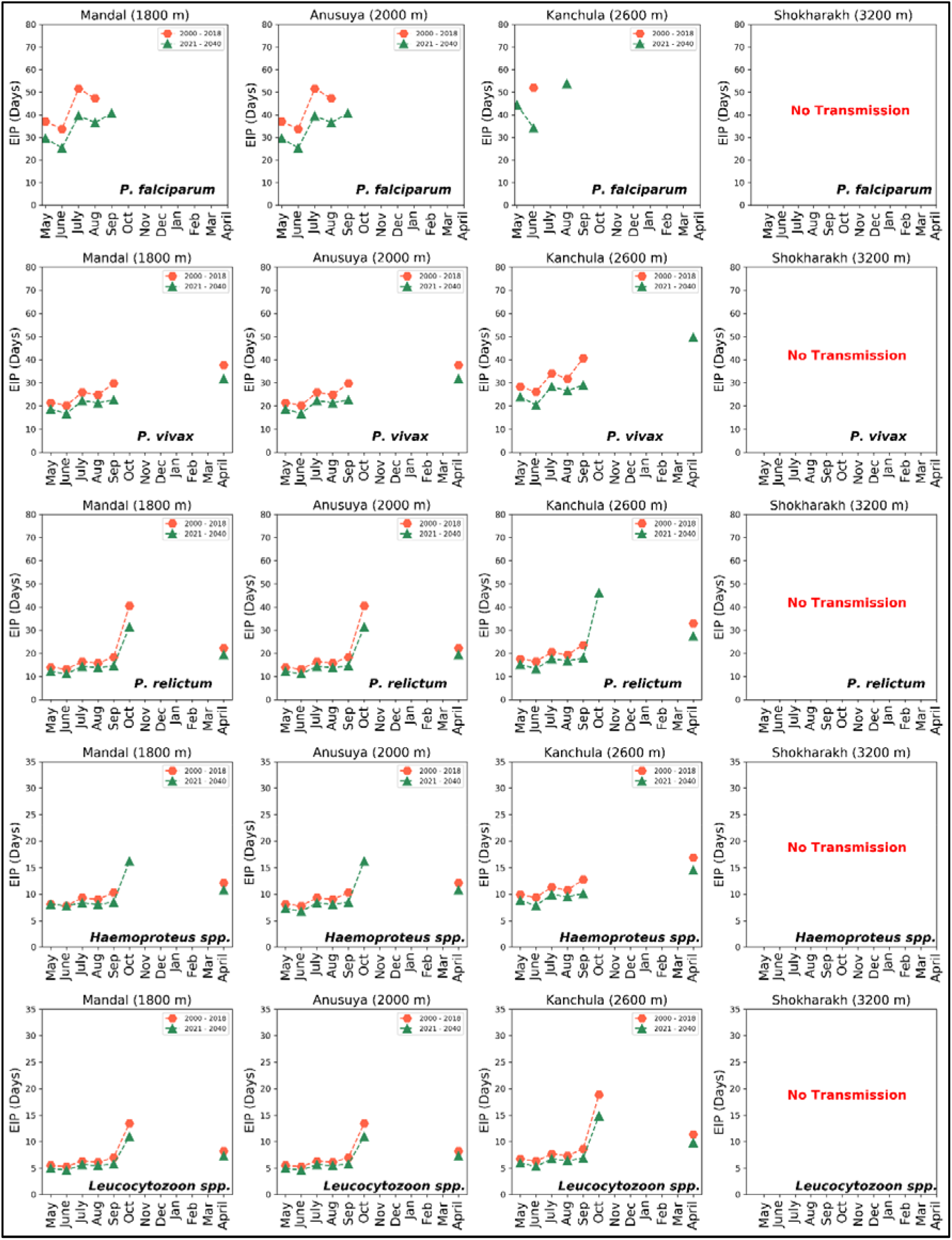
Comparison of extrinsic incubation period (EIP) in days of P. falciparum, P. vivax, P. relictum, Haemoproteus and Leucocytozoon for four sites of the western Himalaya. using mean temperature data (May 2000 to April 2018, hexagon) and future climate data (2021 to 2040, triangle).

## Discussion

Understanding how changing global temperature affects the period during which parasite development can occur is crucial to modelling how the range of viability of the malaria parasite changes with time. We explore these questions using a thermodynamic model that describes the nonlinear relationship between developmental rate and temperature, for five blood-borne parasites and their arthropod hosts. For this we use a parametrization of the EIP proposed by Briére *et al*. (1999), using Bayesian inference methods to estimate the relevant parameters using prior knowledge assembled using an extensive literature search. For avian haemosporidia - *P. relictum, Leucocytozoon* spp., *Haemoproteus* spp. - we are not aware of any prior attempt to obtain the constants defining the EIP as parameterized for the Briére equation. We believe this calculation presents those results for the first time

Using measures of mean temperature versus the diurnal fluctuating temperature in the field, we demonstrated both spatial and temporal variation in malaria transmission risk in the western Himalayan region. Both human malaria and avian malaria prevalence vary with season and intensity across these sites. We showed that for all five vector-borne parasites transmission was most strongly constrained by temperature in high elevation (3200 m) environments throughout the year, and that different models provided largely consistent results for lower elevations.

Whilst the temperature and transmission windows we calculate could fit within the lifespan of *Anopheles* species in general, it is important to note here that our study was primarily designed for the forested habitat with no records of *Anopheles* species (FI unpublished data). Malaria incidence has been reported from human dominated hilly areas below 2000 m with high prevalence for *Anopheles* mosquitoes incriminated as prime malaria vectors (Shukla *et al*. 2007). Our analysis showed that the temperature conditions are not conducive for malaria transmission in the current scenario and the diurnal temperature fluctuation has no effect on malaria transmission biology. In line with the Government of India’s National Framework for Malaria Elimination in India 2016-2030 Program (NVBDCP 2016), it is crucial now to apply such approaches for identification of hotspots using fine-scale data which can help in addressing the ecological drivers of malaria transmission (Mishra *et al*. 2016). These temperature estimates have implications on defining the parasite transmission limits across spatio-temporal scales: i) EIP responds in a non-linear fashion to temperature and is sensitive to small changes in temperature which could have significant effects on the parasite transmission window (e.g., Blanford *et al*. 2013); ii) hourly fluctuations in temperature are experienced in the field by both mosquito and parasite which could provide site-specific insights into parasite transmission range at a small spatial scale. While our data do capture these effects explicitly using mean versus DTR measures, we emphasised that using site-specific data is important for deriving insights into malaria transmission range. Our comparisons of EIP calculated using local meteorological data and WorldClim data showed that relying on weather station data might underestimate the parasite development in a highly seasonal ecosystem with distinct physiographic climatic conditions.

The changing climate has rapidly influenced the rainfall, temperature, and vegetation phenology. These changes are causing shifts in the timing of species activity. For example, a surge in temperature has shifted timing and length of breeding season in birds (Hällfors *et al*. 2020) leading to mismatch with optimal resource abundance which is vital for reproductive success. For short-lived ectotherms, the spread of mosquito species to new habitats in high elevations, short generation times, high population growth rates and strong temperature-imposed selection could lead to fast adaptation (Couper *et al*. 2021).

In the western Himalayan context, the thermodynamic model showed that the EIP days do not support transmission of avian haemosporidians parasites at high elevation sites (3200m). However, EIP days have limited transmission windows for *Plasmodium*, *Haemoproteus* and *Leucocytozoon* from 1800-2000 m. This further implies that low to mid elevation sites have optimal conditions and lack thermal constraints for parasite transmission during peak breeding season (April-May). Ishtiaq and Barve (2018) showed that the probability of infection with *Plasmodium* parasite declines steeply with elevation. In contrast, *Leucocytozoon* spp. infection risk increases with elevation, however, most of these infections were sub microscopic in high elevation in breeding season (April-May). This contrasts with studies on the ecology of haemosporidia in temperate regions where birds with latent infections return to the breeding grounds and experience a relapse, with increase in parasites visible in the blood stages (Applegate 1970, Becker *et al*. 2020). In general, parasite intensity showed a significant decline with elevation in the breeding season (April-May) and an increase across mid-elevations in the nonbreeding season. The absence of gametocytes (infective stage) in the blood during spring season or late emergence of vectors due to environmental conditions could lead to the disruption of transmission cycles (migratory mismatch).

Our data support the hypothesis that avian *Plasmodium*, *Haemoproteus* and *Leucocytozoon* are currently restricted by thermal gradient and provide an ecological explanation for absence of gametocytes in breeding season at high elevation. This further implies that fledgelings might be at the risk of haemosporidian *(Haemoproteus*) infections from June-August which coincides with peak emergence of *Culicoides* spp. in 2000-2600 m sites (Ishtiaq et al. unpublished data). Using a thermodynamic model, we supported the *Plasmodium relictum* transmission scenario in Hawaiian Islands using Degree-day models with 13°C as minimum temperature and 30° C as maximum temperature threshold to complete sporogonic development (LaPointe *et al*. 2010).

We used climate models to interrogate how changing temperature from 2021-2040 could potentially lead to an expansion of the temperature range conducive for malaria transmission in high elevation zones. Using mean temperature data, we found low elevation sites (1800m) might experience unsuitable conditions for parasite transmission in the future which suggests that some habitats that are currently too cool to sustain vector populations may become more favourable in the future, whereas others that are drying may become less conducive to vector reproduction. Therefore, the geographic ranges of mosquitoes may expand or be reduced, which may cause parallel changes in the population of malaria pathogens they transmit. Such expansion also increases the time window of malaria transmission resulting in a larger number of generations of parasites per year that can positively affect parasite abundance (Schroder *et al*. 2008).

One of the limitations of our study is the use of one year data to define these thermal effects at a small spatial scale. To quantify these temperature effects at a fine scale, we need long-term data across multiple sites. Nevertheless, the mean temperature variation using field data and WorldClim records exhibited similar patterns in parasite transmission range. Furthermore, our modelling using mean versus hourly temperatures captures the EIP variation which is corroborated by the prevalence and intensity of parasites characterised in avian hosts. Our data illustrates the contrasting thermal environments that can exist across relatively small spatial scales within a region and can have divergent effects on parasite development.

## Acknowledgements

We thank Wellcome Trust/DBT India Alliance Fellowship (IA/I(S)/12/2/500629) to FI for financial support for this research. We are grateful to Krijn Paaijmans and Lauren Cator for responding to our several queries regarding their work, as well as for providing useful explanations, and Gediminas Valkiũnas for clarifications regarding the range of viable temperatures for *Haemoproteus* and *Leucocytozoon*.

---

**Supplementary Table 1:**
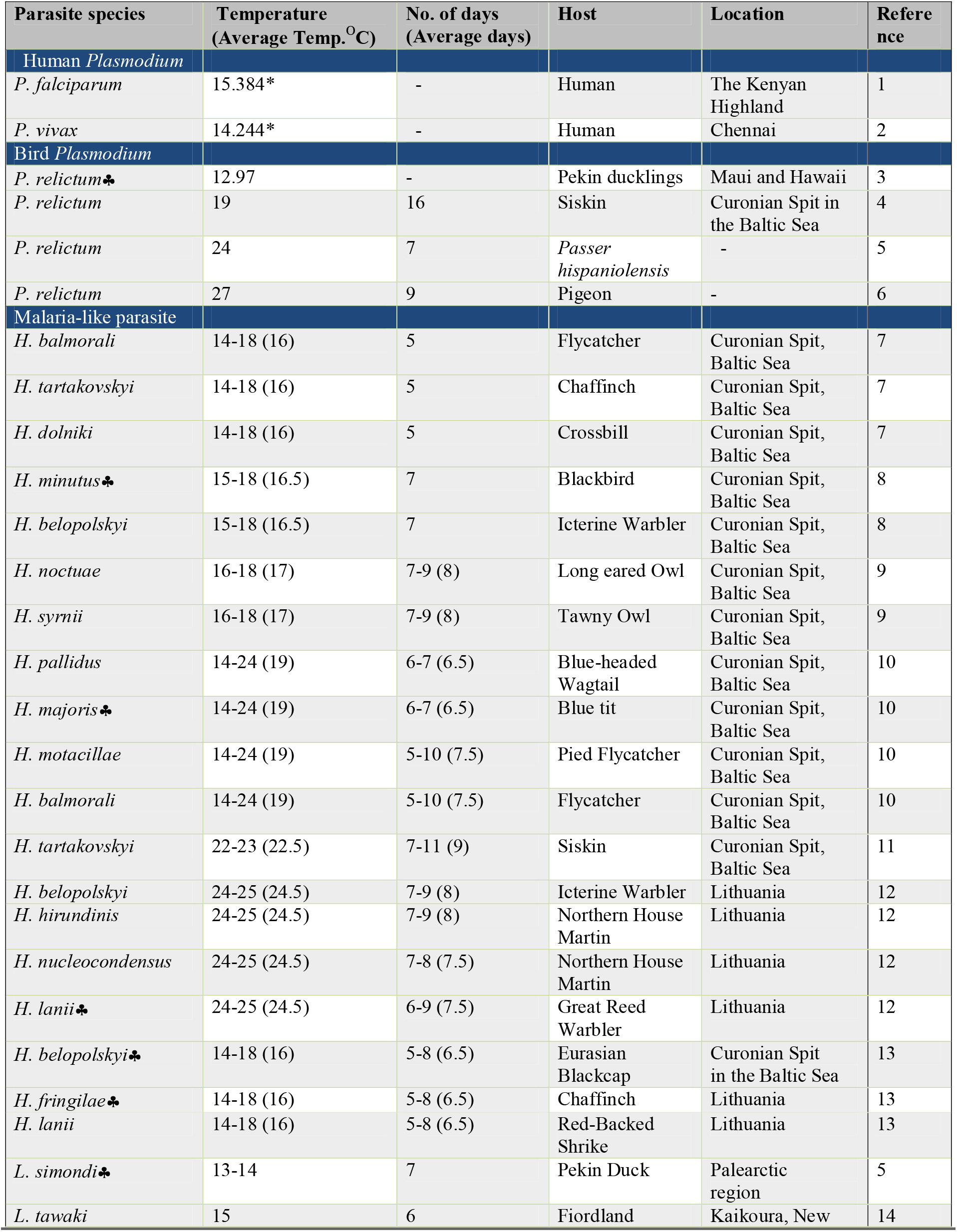
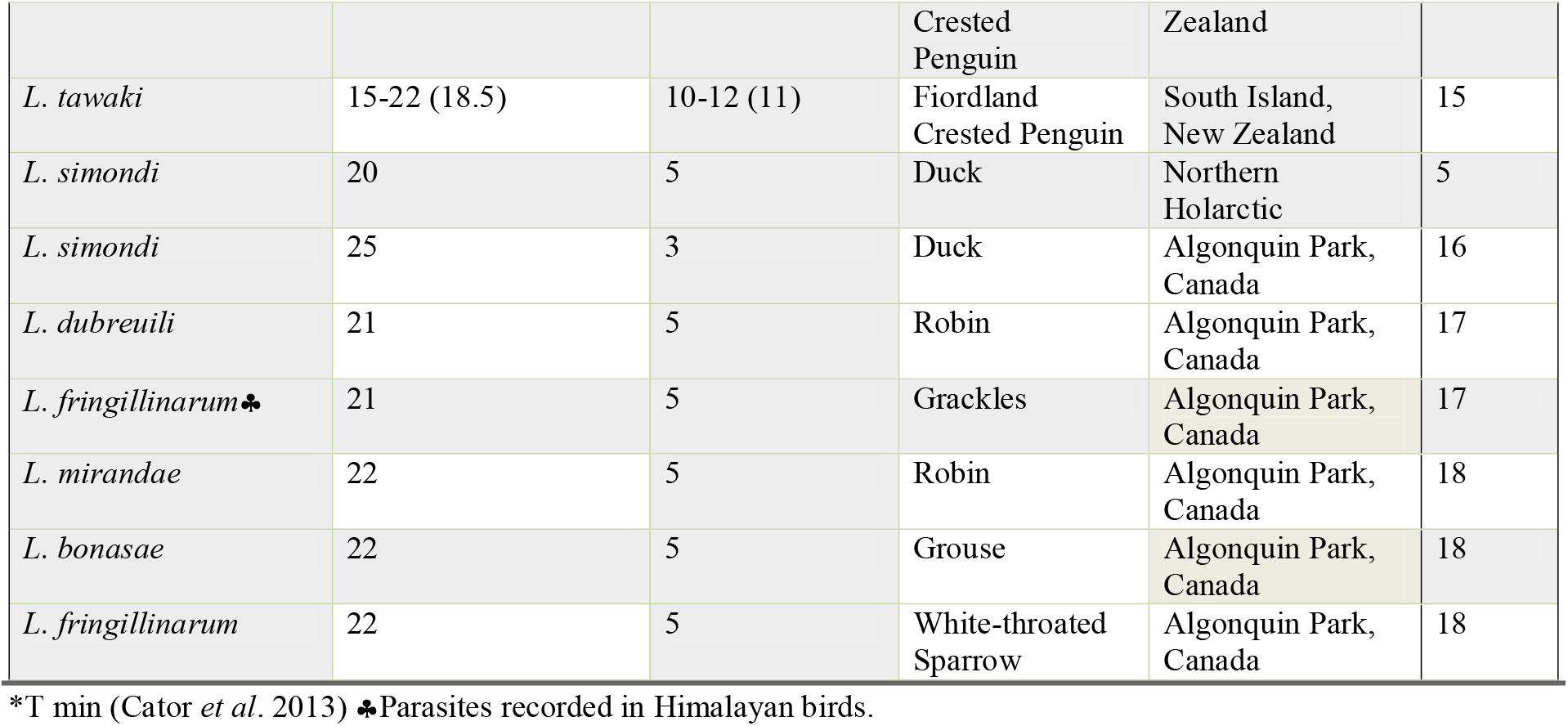
Complete development of *P. falciparum, P. vivax, P. relictum, Haemoproteus* and *Leucocytozoon* in the vector at constant temperatures (°C). No. of days shows total number of days taken to complete the sporogony cycle at constant temperature (°C). Parasite host species is mentioned in the host column. Location refers to where the study has been done.

**Supplementary Table 2:**
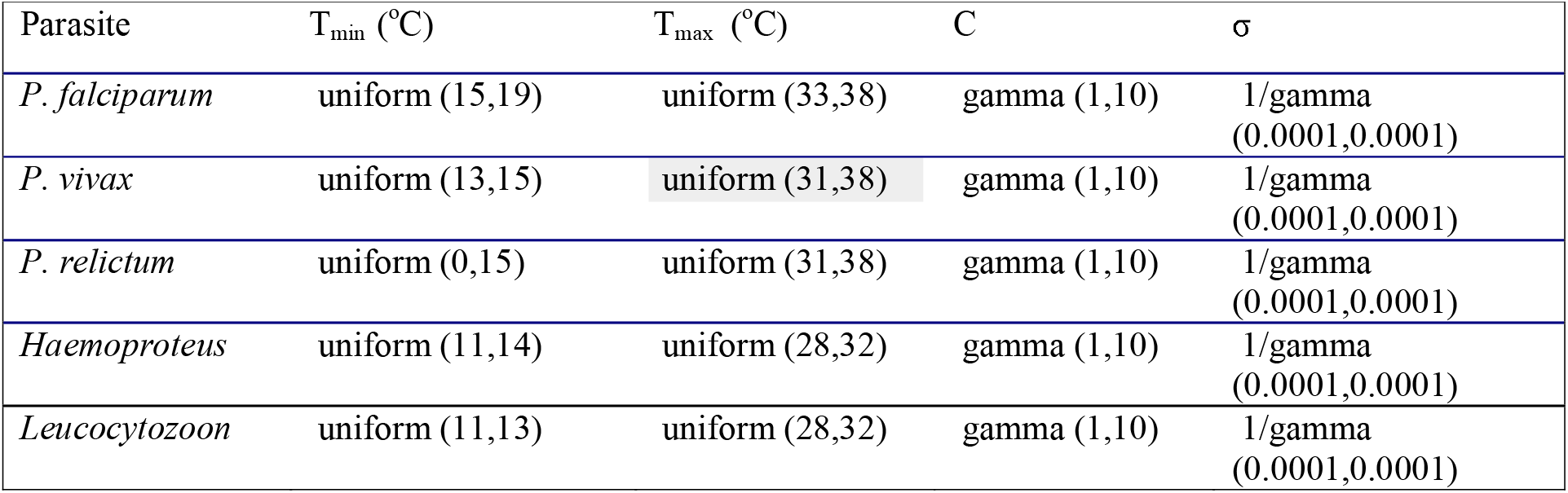
Prior values for the thermodynamic model parameters to estimate using Bayesian modeling

**Figure S1:**
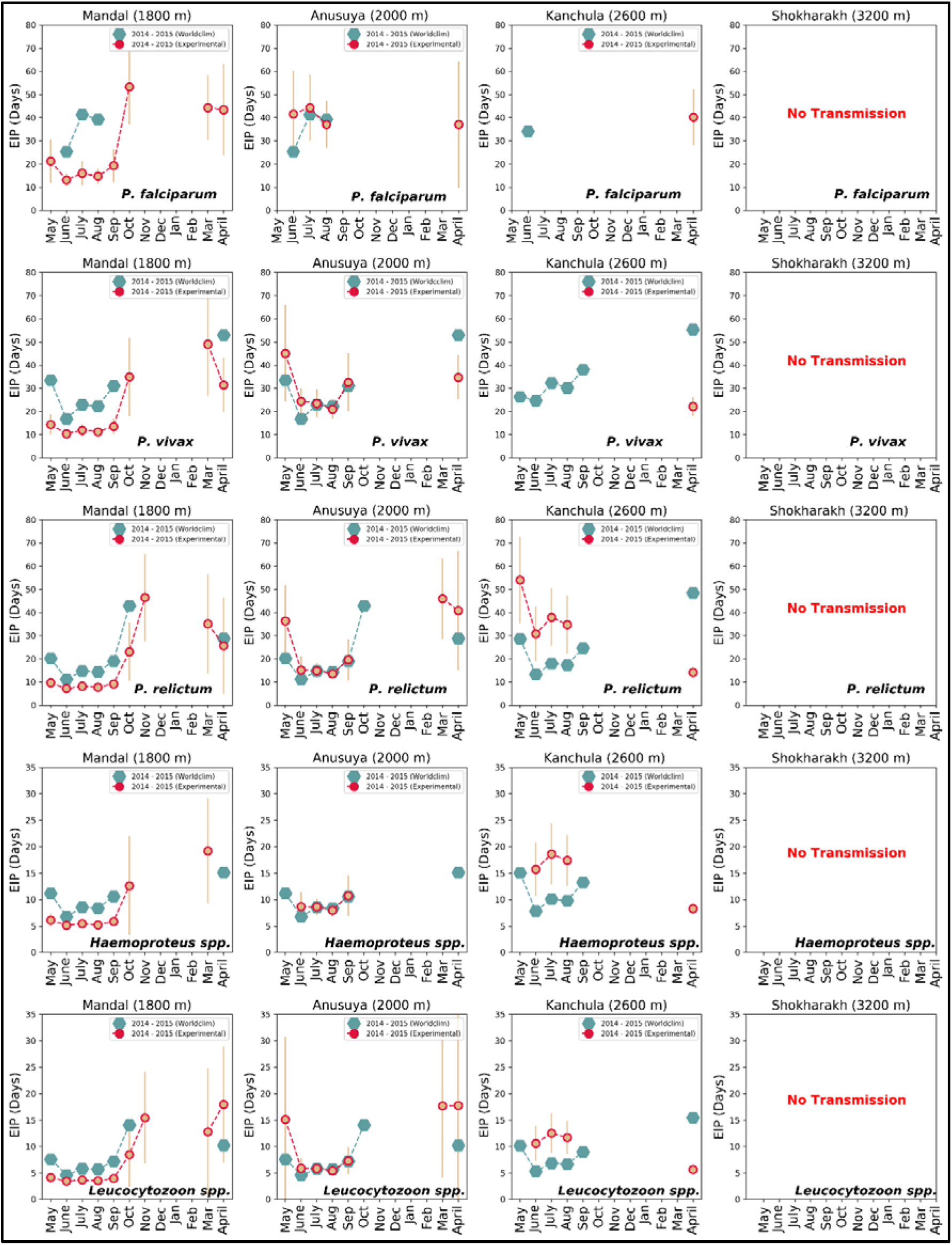
Extrinsic incubation period (EIP) in days of *P. falciparum, P. vivax, P. relictum, Haemoproteus* and *Leucocytozoon* for four sites in the western Himalaya. Calculated using mean temperature gathered from temperature loggers and WorldClim from May 2014 to April 2015. The circle and hexagon show the number of days to complete EIP using mean temperature of the month. Bar represents the standard deviation.

