## Supplementary files for "Exploring the thermal limits of malaria transmission in the western Himalaya"

**Supplementary Table 1:** Complete development of *P. falciparum, P. vivax, P. relictum, Haemoproteus* and *Leucocytozoon* in the vector at constant temperatures (^°^C). No. of days shows total number of days taken to complete the sporogony cycle at constant temperature (^°^C). Parasite host species is mentioned in the host column. Location refers to where the study has been done.

| **Parasite species** | **Temperature**  **(Average Temp. ^O^C)** | **No. of days**  **(Average days)** | **Host** | **Location** | **Reference** |
| --- | --- | --- | --- | --- | --- |
| Human *Plasmodium* |  |  |  |  |  |
| *P. falciparum* | 15.384* | - | Human | The Kenyan Highland | 1 |
| *P. vivax* | 14.244* | - | Human | Chennai | 2 |
| Bird *Plasmodium* |  |  |  |  |  |
| *P. relictum*♣ | 12.97 | - | Pekin ducklings | Maui and Hawaii | 3 |
| *P. relictum* | 19 | 16 | Siskin | Curonian Spit in the Baltic Sea | 4 |
| *P. relictum* | 24 | 7 | *Passer hispaniolensis* | - | 5 |
| *P. relictum* | 27 | 9 | Pigeon | - | 6 |
| Malaria-like parasite |  |  |  |  |  |
| *H. balmorali* | 14-18 (16) | 5 | Flycatcher | Curonian Spit, Baltic Sea | 7 |
| *H. tartakovskyi* | 14-18 (16) | 5 | Chaffinch | Curonian Spit, Baltic Sea | 7 |
| *H. dolniki* | 14-18 (16) | 5 | Crossbill | Curonian Spit, Baltic Sea | 7 |
| *H. minutus*♣ | 15-18 (16.5) | 7 | Blackbird | Curonian Spit, Baltic Sea | 8 |
| *H. belopolskyi* | 15-18 (16.5) | 7 | Icterine Warbler | Curonian Spit, Baltic Sea | 8 |
| *H. noctuae* | 16-18 (17) | 7-9 (8) | Long eared Owl | Curonian Spit, Baltic Sea | 9 |
| *H. syrnii* | 16-18 (17) | 7-9 (8) | Tawny Owl | Curonian Spit, Baltic Sea | 9 |
| *H. pallidus* | 14-24 (19) | 6-7 (6.5) | Blue-headed Wagtail | Curonian Spit, Baltic Sea | 10 |
| *H. majoris*♣ | 14-24 (19) | 6-7 (6.5) | Blue tit | Curonian Spit, Baltic Sea | 10 |
| *H. motacillae* | 14-24 (19) | 5-10 (7.5) | Pied Flycatcher | Curonian Spit, Baltic Sea | 10 |
| *H. balmorali* | 14-24 (19) | 5-10 (7.5) | Flycatcher | Curonian Spit, Baltic Sea | 10 |
| *H. tartakovskyi* | 22-23 (22.5) | 7-11 (9) | Siskin | Curonian Spit, Baltic Sea | 11 |
| *H. belopolskyi* | 24-25 (24.5) | 7-9 (8) | Icterine Warbler | Lithuania | 12 |
| *H. hirundinis* | 24-25 (24.5) | 7-9 (8) | Northern House Martin | Lithuania | 12 |
| *H. nucleocondensus* | 24-25 (24.5) | 7-8 (7.5) | Northern House Martin | Lithuania | 12 |
| *H. lanii*♣ | 24-25 (24.5) | 6-9 (7.5) | Great Reed Warbler | Lithuania | 12 |
| *H. belopolskyi*♣ | 14-18 (16) | 5-8 (6.5) | Eurasian Blackcap | Curonian Spit  in the Baltic Sea | 13 |
| *H. fringilae*♣ | 14-18 (16) | 5-8 (6.5) | Chaffinch | Lithuania | 13 |
| *H. lanii* | 14-18 (16) | 5-8 (6.5) | Red-Backed Shrike | Lithuania | 13 |
| *L. simondi*♣ | 13-14 | 7 | Pekin Duck | Palearctic  region | 5 |
| *L. tawaki* | 15 | 6 | Fiordland Crested  Penguin | Kaikoura, New Zealand | 14 |
| *L. tawaki* | 15-22 (18.5) | 10-12 (11) | Fiordland Crested Penguin | South Island,  New Zealand | 15 |
| *L. simondi* | 20 | 5 | Duck | Northern Holarctic | 5 |
| *L. simondi* | 25 | 3 | Duck | Algonquin Park, Canada | 16 |
| *L. dubreuili* | 21 | 5 | Robin | Algonquin Park, Canada | 17 |
| *L. fringillinarum*♣ | 21 | 5 | Grackles | Algonquin Park, Canada | 17 |
| *L. mirandae* | 22 | 5 | Robin | Algonquin Park, Canada | 18 |
| *L. bonasae* | 22 | 5 | Grouse | Algonquin Park, Canada | 18 |
| *L. fringillinarum* | 22 | 5 | White-throated Sparrow | Algonquin Park, Canada | 18 |

*T min (Cator *et al*. 2013) ♣Parasites recorded in Himalayan birds.

**Supplementary Table 2:** Prior values for the thermodynamic model parameters to estimate using Bayesian modeling

| Parasite | T_min_  (^o^C) | T_max_ (^o^C) | C | σ |
| --- | --- | --- | --- | --- |
| *P. falciparum* | uniform (15,19) | uniform (33,38) | gamma (1,10) | 1/gamma (0.0001,0.0001) |
| *P. vivax* | uniform (13,15) | uniform (31,38) | gamma (1,10) | 1/gamma (0.0001,0.0001) |
| *P. relictum* | uniform (0,15) | uniform (31,38) | gamma (1,10) | 1/gamma (0.0001,0.0001) |
| *Haemoproteus* | uniform (11,14) | uniform (28,32) | gamma (1,10) | 1/gamma (0.0001,0.0001) |
| *Leucocytozoon* | uniform (11,13) | uniform (28,32) | gamma (1,10) | 1/gamma (0.0001,0.0001) |

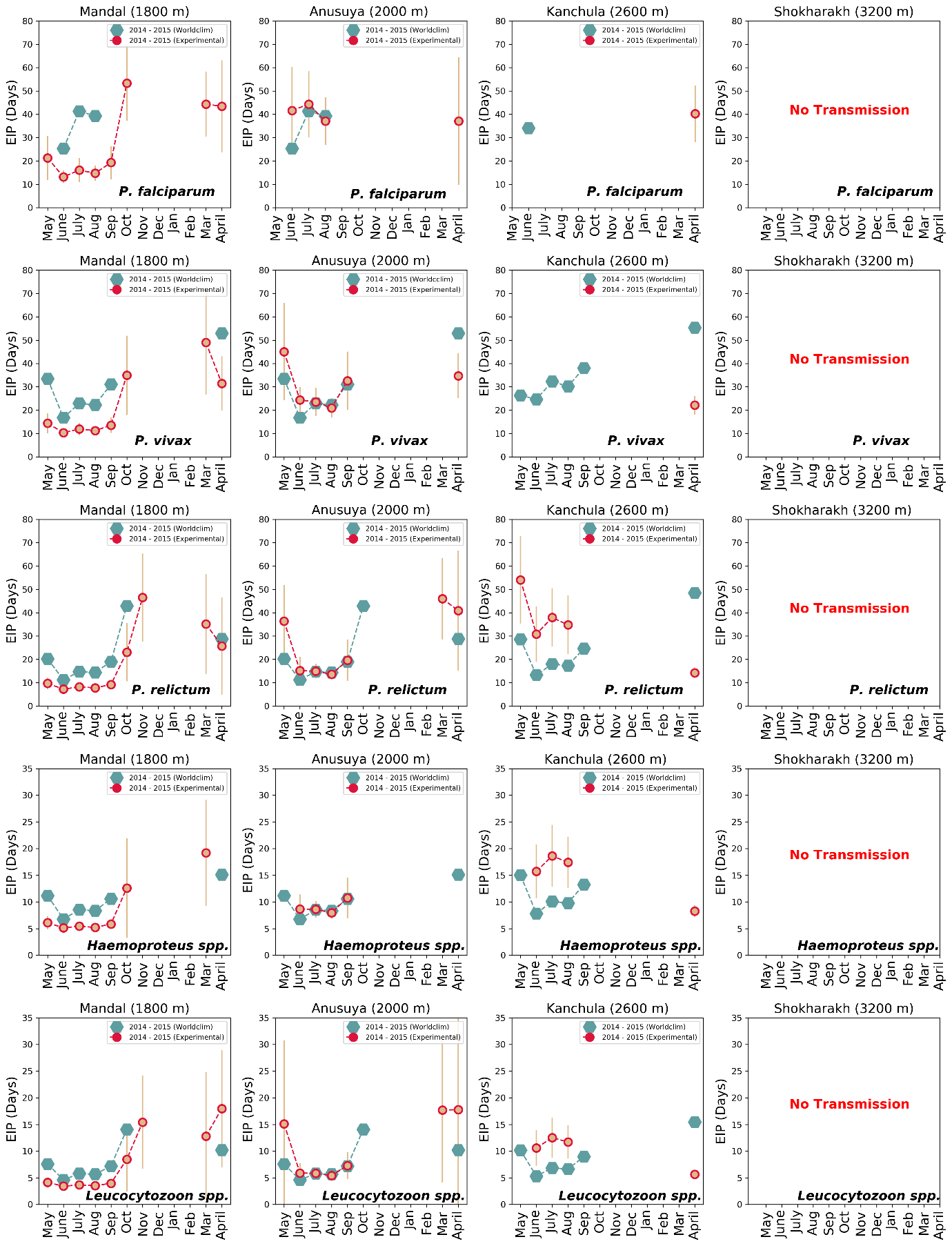

**Figure S1:** Extrinsic incubation period (EIP) in days of *P. falciparum*, *P. vivax*, *P. relictum*, *Haemoproteus* and *Leucocytozoon* for four sites in the western Himalaya. Calculated using mean temperature gathered from temperature loggers and WorldClim from May 2014 to April 2015. The circle and hexagon show the number of days to complete EIP using mean temperature of the month. Bar represents the standard deviation.
